# Genetic architecture of dispersal and local adaptation drives accelerating range expansions

**DOI:** 10.1101/2021.12.02.470932

**Authors:** Jhelam N. Deshpande, Emanuel A. Fronhofer

## Abstract

Contemporary evolution has the potential to significantly alter biotic responses to global change, including range expansion dynamics and biological invasions. However, predictive models often make highly simplifying assumptions about the genetic architecture underlying relevant traits. This can be problematic since genetic architecture defines evolvability, that is, evolutionary rates, and higher order evolutionary processes, which determine whether evolution will be able to keep up with environmental change or not. Therefore, we here study the impact of the genetic architecture of dispersal and local adaptation, two central traits of high relevance for range expansion dynamics, on the speed and variability of range expansions into an environmental gradient, such as temperature. In our theoretical model we assume that dispersal and local adaptation traits result from the products of two non-interacting gene-regulatory networks (GRNs). We compare our model to simpler quantitative genetics models and show that in the GRN model, range expansions are accelerated, faster and more variable. Increased variability implies that these evolutionary changes reduce predictability. We further find that acceleration in the GRN model is primarily driven by an increase in the rate of local adaptation to novel habitats which results from greater sensitivity to mutation (decreased robustness) and increased gene expression. Our results highlight how processes at microscopic scales, here, within genomes, can impact the predictions of large scale, macroscopic phenomena, such as range expansions, by modulating the rate of evolution.

## Introduction

Range expansions and species invasions are happening at an increasing rate as a result of human activities, such as species introductions, the rewiring of dispersal networks [1], and global changes, in general [2]. Therefore, predicting such range expansion and invasion dynamics is of great interest. However, making ecological predictions is challenging because of the various sources of environmental and demographic stochasticity, along with the complexity intrinsic to biological systems. As a consequence, the uncertainty associated with ecological predictions, given a spatial and temporal scale [3], is large, which implies that predictability tends to be small. For example, even in highly controlled laboratory settings, it is not clear if ecological models can correctly predict the speed and uncertainty associated with range expansions [4, 5].

To make matters even worse, contemporary evolution can not only affect ecological patterns and dynamics [6], but also modify the predictability of range expansions [7]. This is because range expansions both drive and are impacted by evolution of traits that influence spatial spread (dispersal ability) and demography (reproduction), forming an eco-evolutionary feedback loop [8]. During range expansions, dispersal evolves due to spatial sorting of dispersers [9, 10], whereby faster dispersers end up at the range front, speeding up range expansions [11]. Therefore, models that include dispersal evolution may better predict range expansions while still globally under-predicting their speed [12]. In addition, range expansion into heterogeneous environments, such as abiotic gradients of temperature, salinity or humidity, may greatly be limited by local adaptation to the environmental gradient [13–15]. However, rapid evolution of local adaptation modifies the demography of the expanding population by changing the survivorship or fecundity in the novel environment. Finally, gene surfing, the spatial analogue of genetic drift [16], introduces stochasticity due to evolution.

While these challenges have been addressed in the literature, relevant models often assume evolution under equilibrium ecological conditions and focus on optimal phenotypes. During range expansions, however, ecological and evolutionary change happens on similar timescales. Therefore, one must not only consider evolutionary optima but also the rate of evolutionary change, that is, evolvability. Evolvability of a trait depends on variation, which refers to the standing genetic variation for the trait, and variability [17], which describes the generation of novel genetic variation by means of mutation or recombination [reviewed in 18]. While variation can be studied using standard quantitative genetics approaches, the study of variability requires knowledge of numbers and effects of loci, their epistatic interaction, pleiotropy, mutation rates, and mutation effects [17]. These properties depend on the genetic architecture, or genotype-to-phenotype map [19–21] of the trait of interest. As a consequence, Kokko et al. [22] argue that adaptive evolution in response to rapid ecological changes is contingent on the supply of genetic variation, which depends on the genetic architecture of underlying traits. Along similar lines, Melián et al. [23] argue that the structure of gene networks has to be taken into account for an appropriate integration of ecology and evolution.

Despite the relevance of a trait’s genetic architecture, eco-evolutionary models of range expansions typically make strong simplifying assumptions regarding the genetic basis of dispersal [24]. Some studies of local adaptation during range expansions have assumed more complex additive [25–27] and non-additive genetic architectures [28, 29]. Other studies have tried to understand the impact of ploidy [30] and modifiers of mutation rates [31] on eco-evolutionary range dynamics. Yet, up to now, models of range expansions into environmental gradients have not taken into account the genetic architecture of both relevant traits: dispersal and local adaptation.

One promising way to integrate genetic architecture into models of range expansions are gene-regulatory network (GRN) models, particularly the one introduced by Wagner [32]. The GRN genotype-to-phenotype (GP) map consists of *n* transcription factors which can regulate each other’s gene expression states (which are the phenotype). The interaction between the genes is represented by a regulatory matrix (the genotype) containing weights. These weights are optimised by evolutionary simulations, where fitness is a function of gene expression levels (phenotype). The non-linearity in the interaction between different loci means that the mutation effects are emergent and not fixed as assumed in quantitative genetics models. Therefore, particularly the robustness (decreased sensitivity) to mutation, also known as genetic canalization, has been studied extensively in the Wagner model and its variants [33–36]. GRN models have also been used to study the evolution of evolvability under conditions of ecological change [37–39], including local adaptation [28, 29, 40].

With this background, the GRN GP map can provide valuable insights into how variation in the direction of adaptation (evolvability) is generated, maintained, and released in relevant traits during range expansions. Here, we develop a novel individual-based model of range expansions into an environmental gradient where dispersal and local adaptation traits can evolve and are assumed to result from the products of two non-interacting gene-regulatory networks (Fig. 1 A). In order to understand the impact of a microscopic property, such as the genetic architecture of a trait, on large-scale, macroscopic processes, here, range expansion dynamics, we compare two alternative models: We either assume that dispersal and local adaptation are encoded by two non-interacting gene-regulatory networks (GRN model), or we assume a model in which one locus each encodes these traits (single locus model).

**Figure 1:**
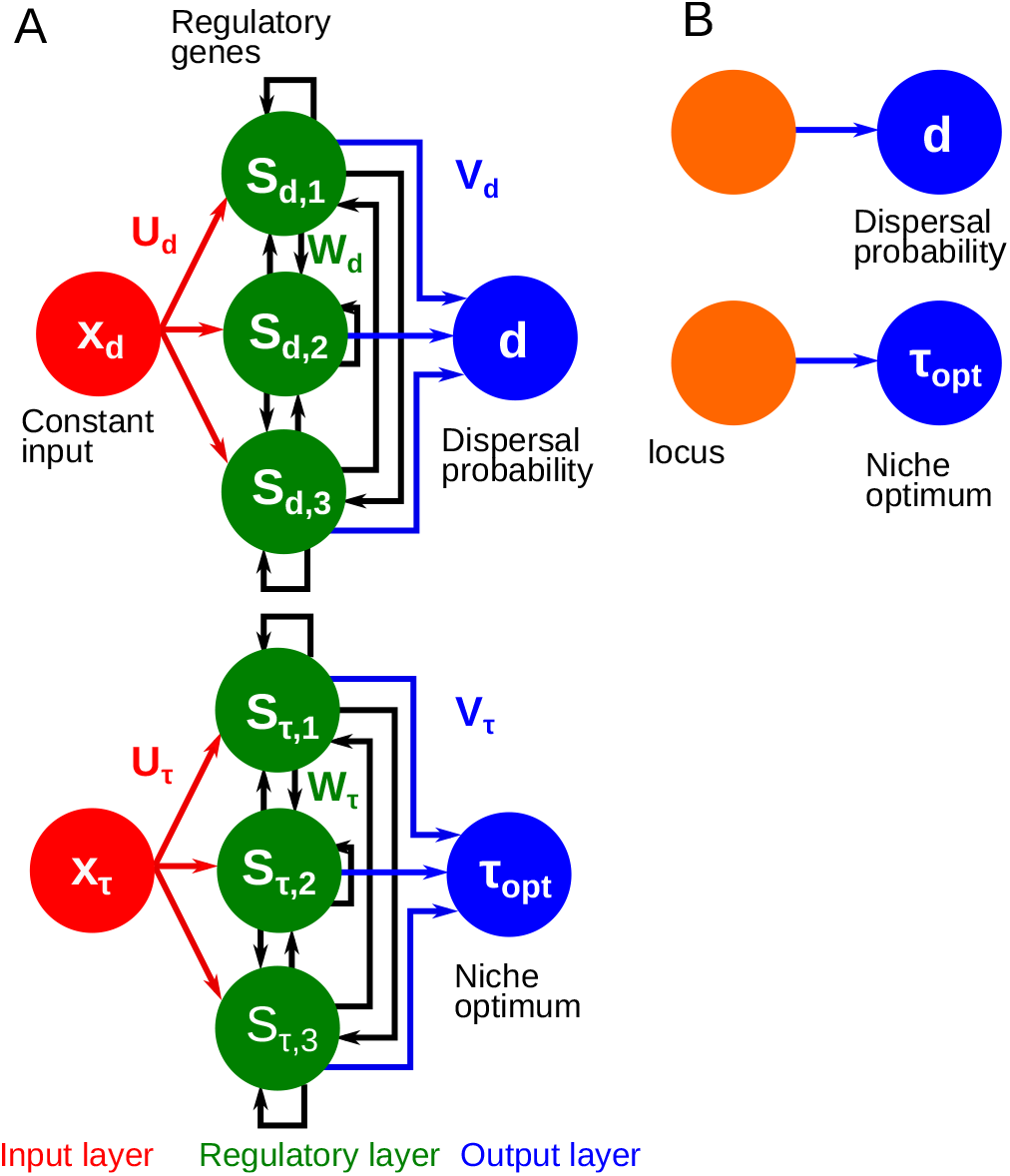
Comparing our model of genetic architecture of traits during range expansions, wherein dispersal and local adaptation traits result from the products of two non-interacting GRNs (GRN model) to a standard quantitative genetics model in which one locus each encodes dispersal and local adaptation (single locus model). A: GRN model. Two different non-interacting GRNs encode dispersal probability (*d*) and the environmental optimum (*τ*_*opt*_) of a Gaussian niche function (the local adaptation trait) separately. Each GRN has an input layer (constant input *x*_*z*_ to each gene, where z represents the dispersal trait *d* or the local adaptation trait *τ*_*opt*_), regulatory layer (regulatory genes with expression level *S*_*z,i*_ for every gene *i* for the trait *z*) and output layer (the phenotype downstream of gene expression, here, dispersal (*d*) or local adaptation (*τ*_*opt*_)). ***U*** _*z*_, ***W*** _*z*_ and ***V*** _*z*_ are matrices of weights that connect the input layer to the regulatory layer, the genes within the regulatory layer and the regulatory genes to the output phenotype respectively for each trait *z*. B: Single locus model. One quantitative locus each encodes dispersal (*d*) and the local adaptation trait (*τ*_*opt*_).

## Methods

### Model overview

We develop an individual-based, discrete-time, discrete-space metapopulation model of range expansions into an external environmental gradient (e.g., climatic gradient). Generations are non-overlapping, with local density-regulation. Dispersal and local adaptation (center of a Gaussian niche) are continuous traits encoded by two non-interacting gene-regulatory networks (GRN) and can evolve.

The landscape is composed of 500 *×* 5 patches. Only the native habitat, that is, the central 10 *×* 5 patches, is occupied for the burn-in period of the first 20000 generations, allowing the traits to reach (quasi)equilibrium. The native habitat has a constant external environment *τ*_0_. After the burn-in period, range expansions begin towards both sides of the landscape into a linear environmental gradient of slope *b*, only in the *x* direction. The landscape can be thought of as a hollow tube extending in the x-direction. Therefore, the boundary conditions in y-direction are periodic which prevents artefacts due to edge-effects. During the burn-in period, the boundary conditions in the x-direction are reflecting. Afterwards, remaining patches on either side become accessible to the individuals, and they can begin range expansion. The simulation ends when the leftmost and rightmost patches are occupied.

After they are born, individuals disperse with a probability *d* which is represented by a GRN, to one of their eight nearest neighbors. During dispersal individuals may die with a probability *µ*, which represents dispersal costs [41]. Individuals mate and reproduce within their target patches. The Beverton and Holt [42] model gives the mean number of offspring each female produces. Local adaptation to the external environment is modelled using a Gaussian niche function with mean *τ*_*opt*_ and niche width *ω*. Therefore, the density-independent survivorship of the offspring is a Gaussian function of the mismatch (level of adaptation *s*) between the local environment *τ* (*x*) and the genetically encoded niche optimum *τ*_*opt*_. Entire patches may go extinct with a probability *ϵ* due to external catastrophic events. The detailed model description and equations can be found in the supplementary information.

### Gene-regulatory network model

The dispersal and local adaptation traits are assumed to result from the effects of two non-interacting gene-regulatory networks (Fig. 1 A). We introduce modifications to the Wagner [32] model similar to Draghi and Whitlock [38] and van Gestel and Weissing [39]. We model the gene-regulatory network as having three layers, the input layer, the regulatory layer and the output layer [39]. The regulatory layer corresponds to Wagner’s model, it contains the genes that characterize the developmental state that is being modeled. These gene expression states (**S**_**I**_) are continuous and sigmoid functions of the input they receive. Each gene receives input from the input layer, which could potentially be an external environmental or an internal cue but in this case we hold it constant for both traits. The input matrix (***U***) connects the input layer to the regulatory layer. The interaction between genes is represented by the regulatory matrix (***W***). Finally, we assume that the trait under consideration varies linearly with gene-expression states. As a consequence, each trait is given by the weighted sum of the gene expression states, and these weights are the output matrix (***V***). Thus, the output layer processes the gene expression states and gives the trait. The gene expression states are iterated for *I* = 20 times for each trait of each individuals, and if a fixed point steady state is reached then the linear combination of these equilibrium gene expression states (**S**^*^) is the phenotype. Limit cycle dynamics are not considered viable [32]. The input matrix, regulatory matrix and output matrix along with the threshold and slope of the sigmoid are genetically encoded by diploid loci. These loci are initialized randomly using a Gaussian distribution with mean 0 and standard deviation 1.

### Analysis

We use two alternate models (Fig. 1), in which dispersal and local adaptation traits are both encoded by two separate GRNs (GRN model) or loci (single locus model). We run 50 replicate simulations which amounts to 100 range expansions and track the range border position as a function of time. For the chosen focal scenario, we run 10 additional replicate simulations or 20 range expansions to follow the dynamics of the level of adaptation (*s*), the sensitivity to mutation (for details, see the supplementary information), and the gene expression states of the local adaptation GRN every 15 patches from the range core. The sensitivity to mutation is a measure of how much a trait changes if the GRN is perturbed.

Additionally, we calculate the ‘adaptation time lag’ which is the difference between the time when the expanding population enters the given patch cross-section and the time when the population adapts to it completely. We use a threshold for the level of adaptation *s >* 0.96, because the population cannot reach *s* = 1 as a consequence of expansion or genetic load. This allows us to track whether the time taken to completely adapt to a given environment (patch-cross section), hence, the rate of adaptation, changes as the expanding population moves farther along the gradient.

## Results and discussion

We find that the GRN model leads to overall faster range expansions, and, most importantly, to accelerating dynamics, when contrasted with the single locus model (Fig. 2). The acceleration observed in the GRN model is robust to changes in model parameters, that is, for environmental gradients of different steepness (*b*), changing dispersal costs [41] (*µ*), and stochastic events such as random patch extinctions (*E*; Fig. S1–S2).

**Figure 2:**
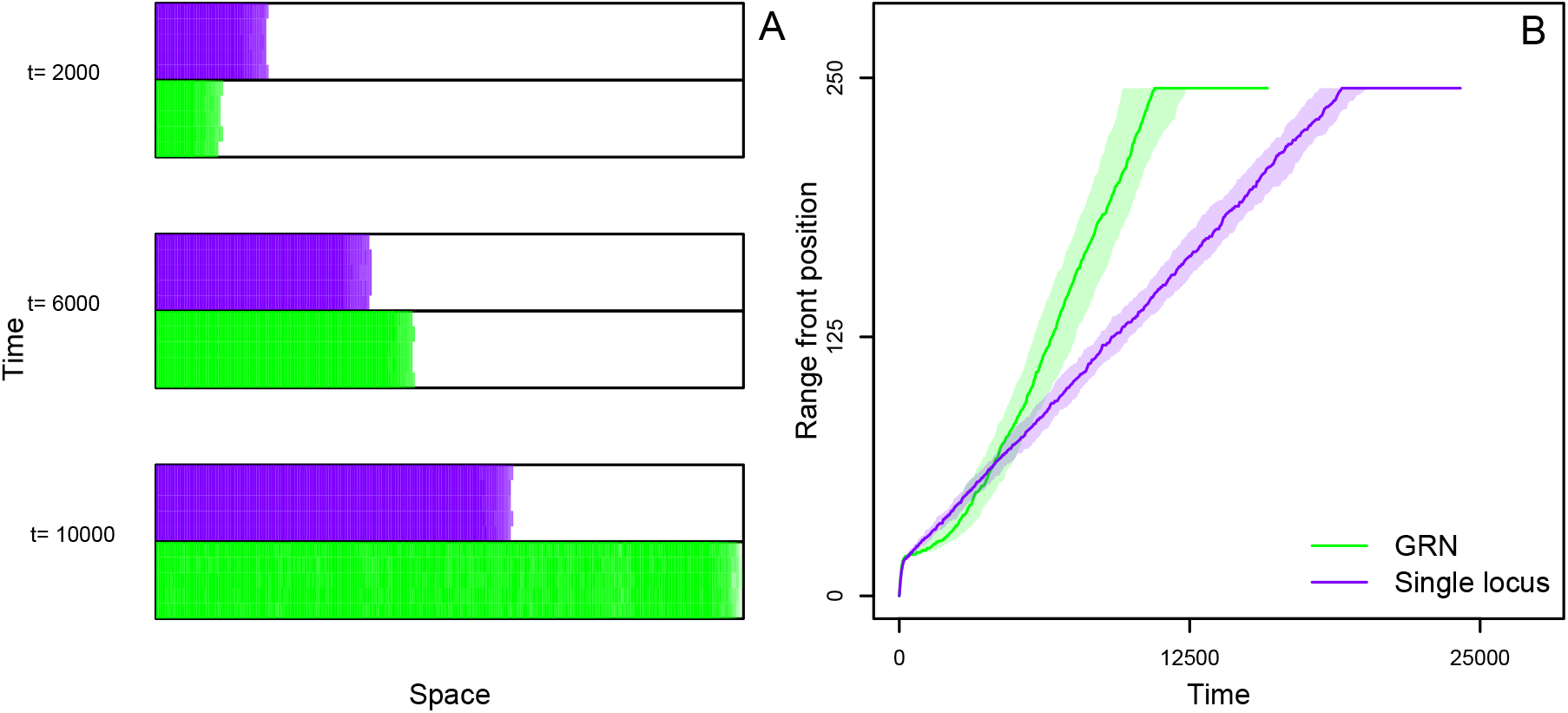
Comparing range dynamics for GRN and single locus genetic architectures. A: Example snapshot of the 250 *×* 5 landscape at different times (*t* = 2000, 6000, 10000) since the beginning of range expansion for single locus (purple) and GRN (green) genetic architectures. Colored patches are occupied, and blank patches are unoccupied. The environmental gradient increases from left to right. Early in the range expansion (*t* = 2000, 6000), new patches are colonized more quickly in the single locus model but accelerating range expansions in the GRN model invert this pattern later on (*t* = 10000). B: Range front position, that is, the position of the occupied patch farthest from the range core, as a function of time since the beginning of range expansion. The solid line represents the median position of the range front over 100 replicate range expansions while the shading represents the corresponding interquartile range. The GRN model leads to overall faster and accelerated range expansions. Focal scenario parameters: slope of gradient *b* = 0.04, fecundity of the females *λ*_0_ = 2, intra-specific competition coefficient *α* = 0.01, mutation rate during range expansions *m*_*min*_ = 0.0001, dispersal cost *µ* = 0.1, extinction probability *E* = 0, niche width *ω* = 1, number of genes per GRN *n* = 3.

Why do we observe accelerated range expansions in the GRN model but not in the single locus model? Fundamentally, rapid evolution of either dispersal or of demographic traits during the range expansion process could modify expansion speeds.

### Dispersal evolution

Previous theoretical and empirical work has shown that range expansions can exhibit accelerating dynamics due to dispersal evolution without considering complex genetic architectures [11, 43]. While we recapture these classical results without an environmental gradient even in a single locus model (Fig. S3), the mechanism of acceleration is not robust to the presence of environmental gradients. In scenarios with environmental gradients where we alter the genetic architecture of dispersal we find that accelerating range expansions are observed only when the local adaptation trait is encoded by a GRN, irrespective of the genetic architecture of dispersal (Fig. S4–S5).

### Evolution of local adaptation

As a consequence, as soon as environmental gradients are present, which is very likely the case in nature [e.g., climatic gradients 44], our model suggests that dispersal evolution may not be responsible for accelerating range expansion dynamics. Therefore, acceleration along gradients must be due to changes in demographic traits. Here, changes in demographic traits are mediated by the evolution of local adaptation which impacts female fecundity and hence demography.

In order to better understand the dynamics of local adaptation, we follow the time a population needs to adapt to a patch across the gradient (averaged over *y* values since there is no gradient in the y-direction). We term the time a population needs to fully adapt (here defined as level of adaptation *s >* 0.96) to a novel environment after a first colonization the ‘adaptation time lag’. The adaptation time lag for the single locus model does not change as the population expands along the gradient (Fig. 3 A). However, in the GRN model, the farther away a patch is from the range core, the smaller the adaptation time lag (Fig. 3 A). This means that, as the population moves along the environmental gradient, it adapts more quickly to new local conditions (for example dynamics of local adaptation in both models see Fig. S6).

**Figure 3:**
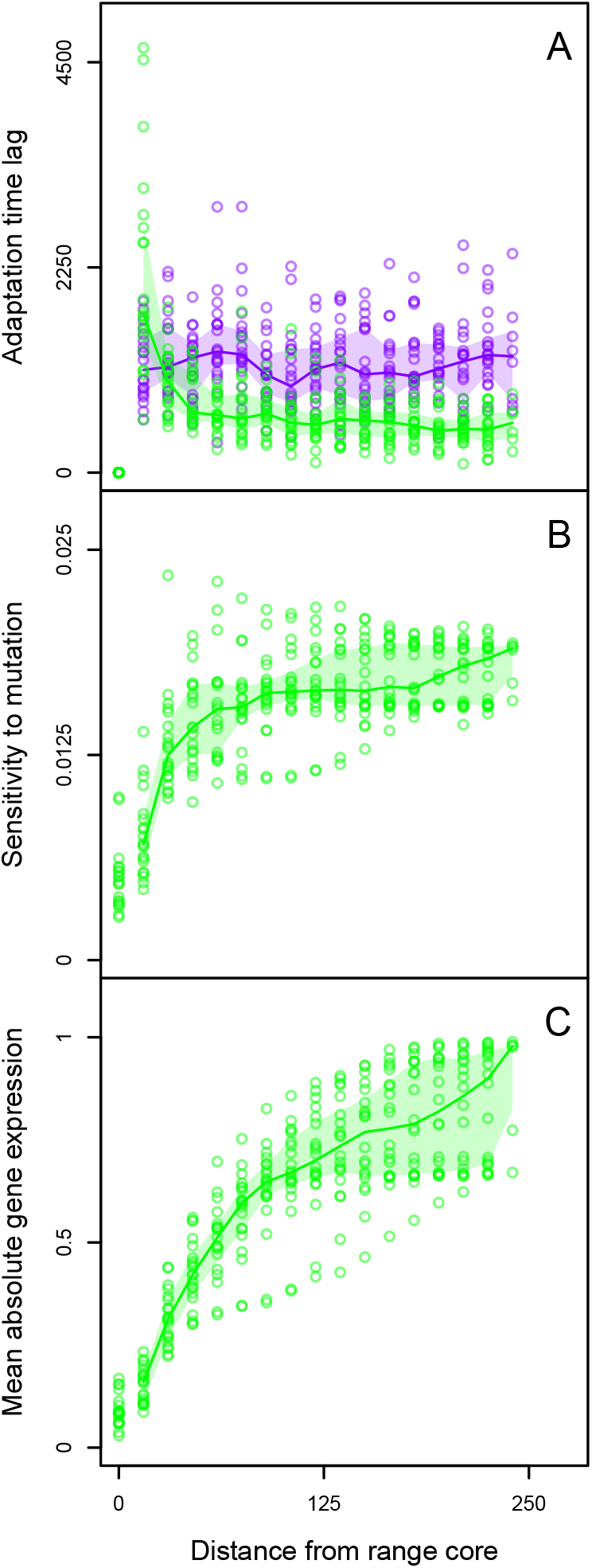
Adaptation time lag, sensitivity to mutation and mean absolute gene expression levels as a function of the distance from the range core. A: The adaptation time lag as a function of the distance from the range core in the GRN model (green) and the single locus model (purple) over 20 replicate range expansions. The adaptation time lag is the difference between the time at which the expanding population completely adapts to a given patch cross-section and when it first arrives there. B: Sensitivity to mutation of the local adaptation trait averaged over the time taken for the expanding population to completely adapt to a given environment as a function of distance from the range core. C: Mean absolute gene expression averaged over the duration of the adaptation time lag as a function of distance from the range core. Focal scenario parameters: slope of gradient *b* = 0.04, fecundity of the females *λ*_0_ = 2, intra-specific competition coefficient *α* = 0.01, mutation rate during range expansions *m*_*min*_ = 0.0001, dispersal cost *µ* = 0.1, extinction probability *E* = 0, niche width *ω* = 1, number of genes per GRN *n* = 3.

Why does the rate of adaptation increase in the GRN model, but remain constant in the single locus model? One crucial difference between the assumptions of the GRN model and the single locus model is how mutation changes the phenotype, that is, the sensitivity to mutation. If a mutation is introduced into one of the alleles encoding local adaptation in the single locus model, the phenotype will change linearly with this mutation. However, in the GRN model, the sensitivity to mutation of the local adaptation trait can itself evolve because of non-linearities in the interaction between the multiple loci. Here, the sensitivity to mutation increases in a saturating manner as the population moves farther along the gradient and away from the range core (Fig. 3 B). Interestingly, this is accompanied by the evolution of extreme gene expression levels (Fig. 3 C). The dynamics of change in the rate of adaptation, sensitivity to mutation, and changes in gene expression levels is presented in an example simulation in Fig. S7.

Furthermore, in populations that are equidistant from the range core, the adaptation time lag is smaller for replicates of expanding populations that are more sensitive to mutation (Fig. S8 A). The sensitivity to mutation of the local adaptation trait is greater, corresponding to extreme gene expression levels (Fig. S8 B).

In summary, for the GRN model, the rate of adaptation increases as the expanding population moves along the gradient because of an increase in the sensitivity to mutation in the local adaptation trait. This is consistent with, for example, Gilbert and Whitlock [26], who show in an additive model that adaptation is faster when larger mutational effects (comparable to mutational sensitivity) are possible. However, our model differs from theirs because the sensitivity to mutation itself changes during range expansion. Clearly, the evolution of extreme gene expression levels during range expansion into a gradient contributes to this increased sensitivity. This is because when the gene expression level for a given gene *S*_*i*_ is closer to the extremes (−1 or 1), any perturbation Δ*V*_*i*_ in the corresponding output weight *V*_*i*_ will be amplified. Closer to intermediate gene expression levels (closer to 0), the perturbation will be silenced. Furthermore, selection towards extreme gene expression levels in more than one gene also increases the overall mutational target. This means that more loci can contribute to the local adaptation trait and can be subject to selection.

These results seem superficially similar to those of Kimbrell and Holt [40] and Kimbrell [28] who find that adaptation to a sink or a gradient of sinks from a source population proceeds due to the breakdown on canalization (increase in sensitivity to mutation) in the local adaptation trait. However, upon closer examination, it is clear that diverging model assumptions lead to differences in the predicted mechanism for increased mutational sensitivity and the resulting adaptation dynamics. These authors assume that gene expression states are the (discrete) phenotype, whereas following Draghi and Whitlock [38], we assume that their downstream effects contribute to a continuous phenotype. When the phenotype under consideration is the gene expression state, then stabilising selection for extremes of gene expression leads to greater genetic canalization [36]. However, in our model, we find that extreme gene expression is associated with greater mutational sensitivity. This is because when a continuous phenotype results from the linear effects of regulatory genes, extremes in gene expression amplify mutations in their downstream effects (output weights). This difference is apparent because in Kimbrell and Holt [40] the breakdown of canalization is only transient, and sensitivity to mutation decreases in the sink as soon as individuals adapt to it. In our model, the sensitivity to mutation does not immediately decrease even after the expanding population has adapted to the new environment (for example, Fig. S7 B). This clearly has consequences for the dynamics of adaptation. Kimbrell and Holt [40] find that adaptation to a patch proceeds through one or more discrete steps. By contrast, in our model, both the increase in sensitivity to mutation and adaptation is gradual (Fig. S7 A, B).

### Predictability of range expansions

As described above, the GRN model leads to faster range expansions than the single locus model and this general pattern is similar for different model parameters and genetic architectures of dispersal (Fig. S9). The speed of range expansion decreases with increasing dispersal costs and extinction probability. The decrease in range expansion speed resulting from greater extinction probability differs from the expectation when only dispersal evolves. This is possibly because while dispersal rates are high, random extinctions at the range front decrease the survivorship of individuals at the range front.

But how does the genetic architecture impact the variation between individual range expansion replicates and, thereby, predictability sensu Melbourne and Hastings [4]? Generally speaking, the GRN model leads not only to faster, but also to more variable range expansions (Fig. S10). This holds true for different slopes, dispersal costs, and extinction probabilities, irrespective of the genetic architecture assumed for the dispersal trait. The greater variability of range expansions in the GRN model, when compared to the single locus model, implies that range expansions are less predictable when the GRN encodes local adaptation. This shows that the assumed genetic architecture not only leads to systematically different predictions but also to different uncertainties.

Therefore, we show that genetic architecture is an important variance generating mechanism sensu Williams et al. [7]. This conclusion has two important implications: 1) in biological systems in which local adaptation is encoded by a GRN, range expansion and invasion dynamics my be even more variable than predicted using classical models 2) Vice versa while range expansions may be intrinsically very variable, using appropriate models based on realistic genetic architectures may at least help us to generate accurate predictions of uncertainty. Accurate estimates of uncertainty are important as they define the range of the forecast horizon [3] as well as risk management strategies [45].

## General discussion

Our study highlights the central role of genetic architecture in determining the speed and variability of range expansions into external environmental gradients. We find that models assuming a GRN genetic architecture for local adaptation lead to faster and accelerated range expansions when compared to models that assume a simpler genetic architecture, irrespective how dispersal is encoded. The acceleration results from a faster rate of adaptation due to an increase in the sensitivity to mutation as the expanding population moves along the gradient.

The increasing rate of adaptation observed in our model is an instance of the evolution of evolvability. Particularly, as defined by Wagner and Altenberg [17], there is a change in trait variability (potential to vary) in the direction of evolutionary change. This is because the increased rate of adaptation results from increased mutational sensitivity. Cobben et al. [31] have demonstrated the evolution of evolvability in individual-based simulations of range expansion when dispersal, local adaptation, and mutation rates evolve. They find that the rate of adaptation at the range front increases because of the evolution of increased mutation rates encoded by a modifier locus. For sexual species, this result can be sensitive to the assumed distribution of mutation effects and level of linkage between the modifier locus and local adaptation locus because the modifier locus evolves by hitch-hiking [46]. However, in our model, there is no need for evolution at a modifier locus. Rather, the increase in the rate of adaptation results from the interaction between genes and their downstream effects.

The consideration of mutational effects is important for understanding the rate of adaptation to a gradient landscape, especially in the absence of standing genetic variation. This has been demonstrated in studies of additive genetic architectures: In Schiffers et al. [25], the authors find that the rate of adaptation is greatest when the mutational effect size matches the steepness of the gradient. Gilbert and Whitlock [26] show that adaptation to a novel habitat patch proceeds primarily by mutations of large effects. By modeling local adaptation as a GRN, we allow for the possibility that the sensitivity to mutation itself can change. As a consequence, whereas in the additive architecture studies described above mutational effects are fixed parameters of the model, in our work, they are emergent properties. Therefore, while we re-emphasize the importance of understanding mutation effects, we add to previous work by showing that the sensitivity to mutation increases as the expanding population moves farther along the gradient, which leads to an increase in the rate of adaptation. This is only possible if the assumption of additivity is relaxed, like in the present study.

Kimbrell and Holt [40] and Kimbrell [28] have shown for a GRN model of local adaptation in a source-sink system, in which gene expression states are the phenotype, that canalization breakdown (increase in sensitivity) is followed by adaptation to a sink or gradient of sinks, with a return to pre-adaptation levels of canalization. In our model, we show that sensitivity to mutation also increases when the trait under consideration is continuous, resulting from linear downstream effects of a GRN, however, by a different mechanism. In continuous, sigmoid Wagner-like GRN models [34] where the gene expression states are the phenotype (including Kimbrell and Holt 40 and Kimbrell 28), Rünneburger and Rouzic [36] show that selection for extremes of gene expression leads to greater canalization for genes under selection. They also show that when few genes are under selection for a non-extreme optimum, then canalization evolves via the unselected genes no longer regulating selected genes. This is because of the sigmoid shape assumed for gene expression. Consider a single gene at equilibrium: If one perturbs any of the inputs it receives (parameters of the sigmoid), closer to intermediate gene expression, this will lead to a change in the expression level, but, at extremes, the gene expression states would remain the same (see Fig. 4 A).

**Figure 4:**
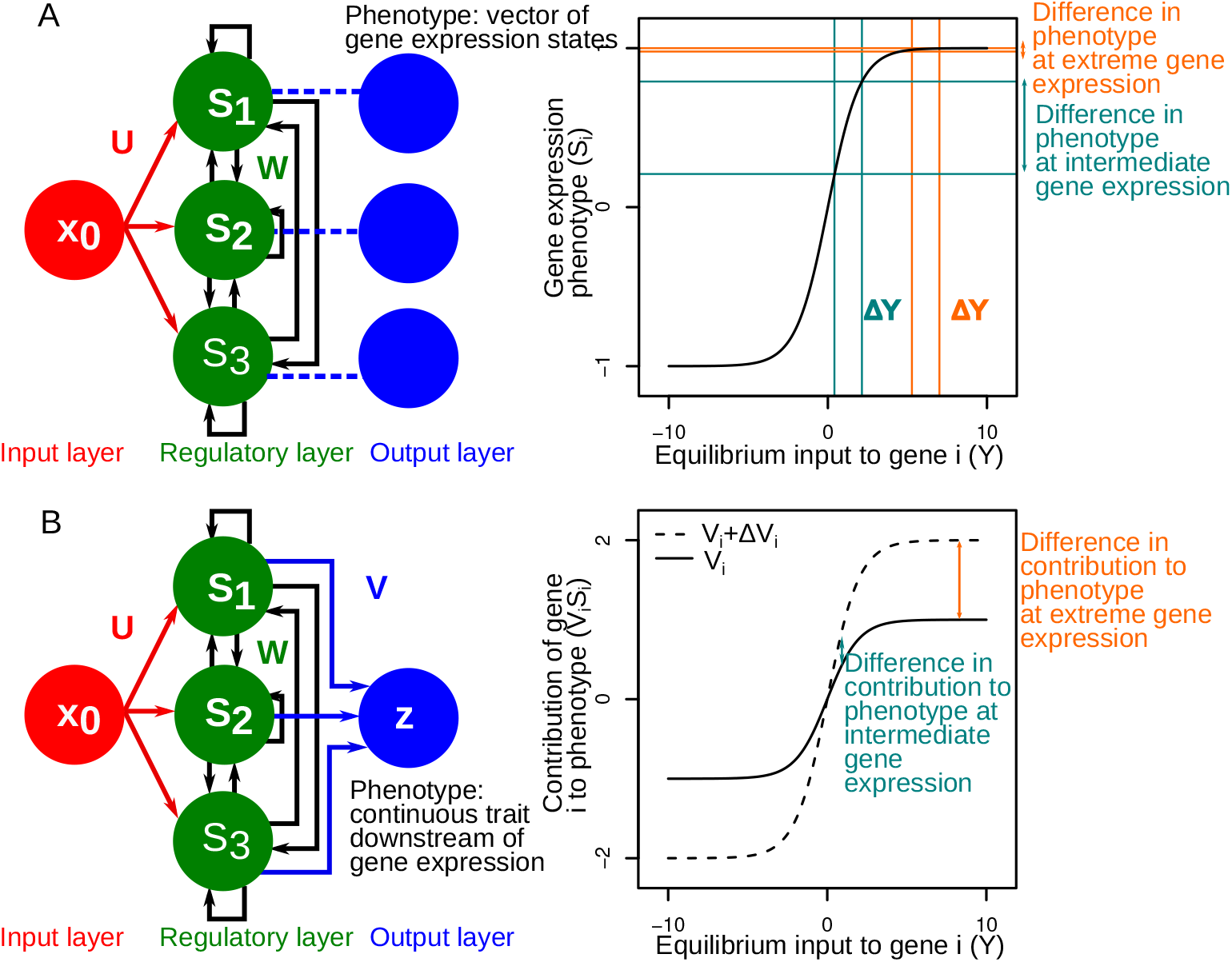
Proposed mechanism for the evolution of increased sensitivity to mutation in the sigmoid Wagner model and the local adaptation GRN in the present study. We have considered a highly simplified scenario in which the GRN is at fixed point equilibrium, and we focus on how a single gene would respond to perturbations to the genotype at extreme and intermediate gene expression levels. We further assume that any perturbation in the input the gene receives takes its gene expression to another fixed point equilibrium. A: Sigmoid Wagner model: In this model, the vector of gene expression states **S** is the phenotype, therefore any perturbation Δ*Y* in the input a single gene receives (either from the input layer or from other genes), would lead to greater phenotypic difference at intermediate gene expression states when compared to those at extreme [36]. B: Our model in which the continuous local adaptation phenotype results from the effects of gene expression. Here, the linear effects downstream of a gene *i* contribute to a trait determined by the output weight *V*_*i*_, therefore the contribution of each gene to the phenotype is *V*_*i*_*S*_*i*_. In this case, any perturbation in the output weights *V*_*i*_, say *V*_*i*_ + Δ*V*_*i*_ would yield a greater difference in contribution to the phenotype at extreme rather than intermediate gene expression.

However, in our model, we find that greater mutational sensitivity is associated with extremes of gene expression when the continuous trait being modelled results from linear downstream effects of regulatory genes. This is because, at extreme gene expression, any perturbation in the downstream effect of the gene (output weight) is amplified (see Fig. 4 B). To our knowledge, this mechanism has been discussed once before by Draghi and Whitlock [47] but with a different kind of GRN model in the context of the evolution of robustness to intrinsic noise. These authors study a trade-off between the robustness of a single gene and a GRN to intrinsic noise (which is greater at low gene expression) and the amplification of mutations in downstream mutation effects (greater at high gene expression) and find that robustness to intrinsic noise evolves in spite of this constraint. The amplification of downstream mutations by extreme gene expression can be considered as a special case of positive directional epistasis, in which multiple loci interact to amplify each other’s effects in the direction of selection. This is a mechanism that has been shown to lead to greater evolvability by Carter et al. [48] in a multilinear GP map [49]. Therefore, based on our study, we hypothesize for a continuous local adaptation trait that one would expect greater sensitivity to mutation farther away from the range core and faster rates of adaptation.

More importantly, we show that genetic architecture has a very consistent impact on the predictability of range expansions. Between-replicate variation is higher when local adaptation and dispersal are encoded by a GRN than by one locus each. Therefore, range expansions are less predictable in the case of the GRN model than the single locus model. This could possibly be because, in the GRN model, uncertainty is propagated between the process of increase in sensitivity to mutation (evolvability) and the adaptation to the new environmental conditions (evolution). The challenges of predicting biological responses to environmental change, in general, are compounded by the levels of organization and spatiotemporal scales [3]. In particular, previous purely ecological studies of range expansion have found that stochastic models may either accurately estimate [5] or overestimate [4] the predictability of observed range expansion speeds. Evolution clearly plays a role in the predictability of range expansion [7]. Going beyond Williams et al. [7], we show that the predictability of range expansions is not only modified by evolution but also the evolution of evolvability. This highlights how assumptions about a lower level of biological organization, here, the assumed genetic architecture, can propagate in a way that may impact not only mean predictions of ecological processes [23], but also their predictability.

## Author contributions

J.N.D. and E.A.F. conceived the study. J.N.D. developed and analysed the simulation model. J.N.D. and E.A.F. interpreted the data. J.N.D. and E.A.F. wrote the manuscript.

## Acknowledgements

This work was supported by a grant from the Agence Nationale de la Recherche (No.: ANR-19-CE02-0015) to EAF. This is publication ISEM-YYYY-XXX of the Institut des Sciences de l’Evolution – Montpellier.

## Data availability

Computer code is available on GitHub: doi: 10.5281/zenodo.5747752

## References

[1] Bullock, J. M., Bonte, D., Pufal, G., da Silva Carvalho, C., Chapman, D. S., García, C., García, D., Matthysen, E., & Delgado, M. M. (2018) Trends Ecol. Evol. 33, 958–970.

[2] Chen, I.-C., Hill, J. K., Ohlemuller, R., Roy, D. B., & Thomas, C. D. (2011) Science 333, 1024–1026.

[3] Petchey, O. L., Pontarp, M., Massie, T. M., Kéfi, S., Ozgul, A., Weilenmann, M., Palamara, G. M., Altermatt, F., Matthews, B., Levine, J. M., Childs, D. Z., McGill, B. J., Schaepman, M. E., Schmid, B., Spaak, P., Beckerman, A. P., Pennekamp, F., & Pearse, I. S. (2015) Ecol. Lett. 18, 597–611.

[4] Melbourne, B. A. & Hastings, A. (2009) Science 325, 1536–1539.

[5] Giometto, A., Rinaldo, A., Carrara, F., & Altermatt, F. (2014) Proc. Natl. Acad. Sci. 111, 297–301.

[6] Govaert, L., Fronhofer, E. A., Lion, S., Eizaguirre, C., Bonte, D., Egas, M., Hendry, A. P., De Brito Martins, A., Melián, C. J., Raeymaekers, J. A. M., Ratikainen, I. I., Saether, B.-E., Schweitzer, J. A., & Matthews, B. (2019) Funct. Ecol. 33, 13–30.

[7] Williams, J. L., Hufbauer, R. A., & Miller, T. E. X. (2019) Trends Ecol. Evol. 34, 903–913.

[8] Miller, T. E., Angert, A. L., Brown, C. D., Lee-Yaw, J. A., Lewis, M., Lutscher, F., Marculis, N. G., Melbourne, B. A., Shaw, A. K., Szuűcs, M., et al. (2020) Ecology 101, e03139.

[9] Shine, R., Brown, G. P., & Phillips, B. L. (2011) Proc. Natl. Acad. Sci. 108, 5708–5711.

[10] Phillips, B. L. & Perkins, T. A. (2019) Theor. Ecol. 12, 155–163.

[11] Phillips, B. L., Brown, G. P., Webb, J. K., & Shine, R. (2006) Nature 439, 803–803.

[12] Perkins, T. A., Phillips, B. L., Baskett, M. L., & Hastings, A. (2013) Ecol. Lett. 16, 1079–1087.

[13] Kirkpatrick, M. & Barton, N. (1997) Am. Nat. 150, 1–23.

[14] García-Ramos, G. & Rodríguez, D. (2002) Evolution 56, 661.

[15] Szuűcs, M., Vahsen, M. L., Melbourne, B. A., Hoover, C., Weiss-Lehman, C., & Hufbauer, R. A. (2017) Proc. Natl. Acad. Sci. 114, 13501–13506.

[16] Slatkin, M. & Excoffier, L. (2012) Genetics 191, 171–181.

[17] Wagner, G. P. & Altenberg, L. (1996) Evolution 50, 967.

[18] Pigliucci, M. (2008) Nat. Rev. Genet. 9, 75–82.

[19] Alberch, P. (1991) Genetica 84, 5–11.

[20] Pigliucci, M. (2010) Philos. Trans. R. Soc. B-Biol. Sci. 365, 557–566.

[21] Nichol, D., Robertson-Tessi, M., Anderson, A. R. A., & Jeavons, P. (2019) J. R. Soc. Interface 16, 20190332.

[22] Kokko, H., Chaturvedi, A., Croll, D., Fischer, M. C., Guillaume, F., Karrenberg, S., Kerr, B., Rolshausen, G., & Stapley, J. (2017) Trends Ecol. Evol. 32, 187–197.

[23] Melián, C. J., Matthews, B., de Andreazzi, C. S., Rodríguez, J. P., Harmon, L. J., & Fortuna, M. A. (2018) Trends Ecol. Evol. 33, 504–512.

[24] Saastamoinen, M., Bocedi, G., Cote, J., Legrand, D., Guillaume, F., Wheat, C. W., Fronhofer, E. A., Garcia, C., Henry, R., Husby, A., et al. (2018) Biol. Rev. 93, 574–599.

[25] Schiffers, K., Schurr, F. M., Travis, J. M. J., Duputié, A., Eckhart, V. M., Lavergne, S., McInerny, G., Moore, K. A., Pearman, P. B., Thuiller, W., Wuüest, R. O., & Holt, R. D. (2014) Ecography 37, 1218–1229.

[26] Gilbert, K. J. & Whitlock, M. C. (2017) J. Evol. Biol. 30, 591–602.

[27] Gilbert, K. J., Sharp, N. P., Angert, A. L., Conte, G. L., Draghi, J. A., Guillaume, F., Hargreaves, L., Matthey-Doret, R., & Whitlock, M. C. (2017) Am. Nat. 189, 368–380.

[28] Kimbrell, T. (2010) Evol. Ecol. 24, 891–909.

[29] Malcom, J. W. (2011) PLoS ONE 6, e21541.

[30] Kubisch, A., Holt, R. D., Poethke, H.-J., & Fronhofer, E. A. (2014) Oikos 123, 5–22.

[31] Cobben, M. M. P., Mitesser, O., & Kubisch, A. (2017) BMC Evol. Biol. 17.

[32] Wagner, A. (1994) Proc. Natl. Acad. Sci. 91, 4387–4391.

[33] Wagner, A. (1996) Evolution 50, 1008.

[34] Siegal, M. L. & Bergman, A. (2002) Proc. Natl. Acad. Sci. 99, 10528–10532.

[35] Ciliberti, S., Martin, O. C., & Wagner, A. (2007) Proc. Natl. Acad. Sci. 104, 13591–13596.

[36] Rünneburger, E. & Rouzic, A. L. (2016) BMC Evol. Biol. 16.

[37] Draghi, J. & Wagner, G. P. (2009) J. Evol. Biol. 22, 599–611.

[38] Draghi, J. A. & Whitlock, M. C. (2012) Evolution 66, 2891–2902.

[39] van Gestel, J. & Weissing, F. J. (2016) Sci. Rep. 6.

[40] Kimbrell, T. & Holt, R. (2007) Am. Nat. 169, 370–382.

[41] Bonte, D., Van Dyck, H., Bullock, J. M., Coulon, A., Delgado, M., Gibbs, M., Lehouck, V., Matthysen, E., Mustin, K., Saastamoinen, M., Schtickzelle, N., Stevens, V. M., Vandewoestijne, S., Baguette, M., Barton, K., Benton, T. G., Chaput-Bardy, A., Clobert, J., Dytham, C., Hovestadt, T., Meier, C. M., Palmer, S. C. F., Turlure, C., & Travis, J. M. J. (2012) Biol. Rev. Biol. Proc. Camb. Philos. Soc. 87, 290–312.

[42] Beverton, R. J. H. & Holt, S. J. (1957) On the dynamics of exploited fish populations (Chapman & Hall, London).

[43] Kubisch, A., Fronhofer, E. A., Poethke, H. J., & Hovestadt, T. (2013) Am. Nat. 181, 700–706.

[44] Colautti, R. I. & Barrett, S. C. H. (2013) Science 342, 364–366.

[45] Travis, J. M. J., Harris, C. M., Park, K. J., & Bullock, J. M. (2011) Methods Ecol. Evol. 2, 477–488.

[46] Romero-Mujalli, D., Jeltsch, F., & Tiedemann, R. (2019) BMC Evol. Biol. 19.

[47] Draghi, J. & Whitlock, M. (2015) Evolution 69, 2345–2358.

[48] Carter, A. J. R., Hermisson, J., & Hansen, T. F. (2005) Theor. Popul. Biol. 68, 179–196.

[49] Hansen, T. F. & Wagner, G. P. (2001) Theor. Popul. Biol. 59, 61–86.

