## Supplementary information for "Genetic architecture of dispersal and local adaptation drives accelerating range expansions"

### Supplementary Methods

#### Life cycle

After birth, individuals first have the possibility to disperse, following which they mate and reproduce. At each generation, entire patches can go extinct with a probability $\epsilon$, representing temporally and spatially random catastrophic extinction events. Every patch in the landscape is initialised with $N_0 = 100$ individuals which have equal chance of being male or female.

##### Dispersal

Dispersal is natal, and individuals can disperse to one of the eight surrounding patches (nearest-neighbour eight dispersal) with a probability $d$, determined by its genetic architecture (described below). Dispersing individuals may die with a probability $\mu$ while dispersing, which captures the multiple costs of dispersal (1).

##### Reproduction and inheritance

After dispersal, individuals mate in their destination patches. Each female mates with a male drawn randomly with replacement from the local population, reflecting female choice. The population dynamics follow logistic growth according to the Beverton-Holt model (2) with local density regulation:

$$N_{x,y,t+1} = N_{x,y,t} \frac{\lambda_0}{1 + \alpha N_{x,y,t}} \tag{S1}$$

where $\lambda_0$ is the intrinsic growth rate, and $\alpha$ is the intraspecific competition coefficient. $K = \frac{\lambda_0 - 1}{\alpha}$ is the expected equilibrium density in the Beverton-Holt model. Each female produces a number of offspring drawn from a Poisson distribution with a mean given by $\lambda = \frac{2\lambda_0}{1+\alpha N_{x,y,t}}$, which models demographic stochasticity. The factor of two accounts for the fact that only half of the population (females) are capable of bearing offspring and makes $\lambda$ interpretable at the population level. The offspring have an equal probability of being male or female, therefore the expected sex ratio is 0.5. After reproduction, the adult individuals die and are replaced in the metapopulation by their offspring. The modifications to this model to understand local adaptation will be explained in the following section.

Each parameter of a dispersal genetic architecture is represented by a locus, with two alleles. Offspring inherit one allele from the mother and one from the father, randomly. The alleles may be modified with a mutation rate $m(t)$ by adding numbers drawn from a normal distribution with mean zero and variance $\sigma_m$. The mutation rate varies with time to generate variation at the beginning of the simulation while allowing the genotypes to stabilise towards the end. Hence, $m(t)$ decreases linearly till the $5000^{th}$ generation and then remains constant (see 3). We chose this approach to allow the genetic algorithm to explore the entire fitness landscape more easily without potentially getting stuck in local optima. Note that mutation rates only vary at the very beginning of the burn-in phase and are constant thereafter. This includes the range expansion phase, implying that any pattern observed during range expansion cannot be due to this mutation procedure choice.

$$m(t) = \begin{cases} m_{max} - \frac{(m_{max}-m_{min})t}{5000} & t < 5000 \\ m_{min} & t \geq 5000 \end{cases} \tag{S2}$$

### Modelling local adaptation

#### Landscape

The equation describing the environmental gradient ($\tau(x)$) as a function of the patch identity $x$ is specified by two parameters: $\tau_0$ is the constant environment in the burn-in region (between $x_{left}$ and $x_{right}$) and $b$ is the slope of the gradient.

$$\tau(x) = \begin{cases} b(x_{left} - x) + \tau_0 & x < x_{left} \\ \tau_0 & x_{left} \leq x \leq x_{right} \\ b(x - x_{right}) + \tau_0 & x > x_{right} \end{cases} \tag{S3}$$

There is no gradient in the y-direction.

#### Local adaptation

The fecundity of a female (or the survivorship of the offspring) when the environment is not optimal is given by a Gaussian niche function. The environmental optimum ($\tau_{opt}$) is genetically encoded (either by a GRN or a single locus, depending on the assumed genetic architecture), and the niche width ($\omega$) is assumed to be fixed. While the male mate does not directly influence $\tau_{opt}$, $\tau_{opt}$ is determined by both the female's mother and father. Therefore, overall there is selection on male and female genotypes. If $1 - s$ is the reduction in fecundity of a female, then:

$$s = \exp \frac{-(\tau(x) - \tau_{opt})^2}{\omega} \tag{S4}$$

Therefore, the number of offspring produced by a female (based on Eq. (S1)) will now be drawn from a Poisson distribution with mean $\lambda = s\frac{2\lambda_0}{1+\alpha N_{x,y,t}}$. Hence, a greater $s$ implies a higher level of adaptation to the external environment of a patch.

### Gene-regulatory network model

#### Background on genotype-to-phenotype maps and gene-regulatory networks

One way in which the genetic architecture of traits has been conceptualised is the genotype-to-phenotype map (GP map). Alberch (4) proposed that the relationship between genotype and phenotype is non-linear and is mapped by dynamic developmental processes, determined by parameters (influenced by the genotype) associated with the developmental system. Stable phenotypes then correspond to steady states of a developmental mapping. According to this conceptualisation, areas in parameter space correspond to phenotypes separated from each other by a boundary. Phenotypes are more robust if they are within a larger region of parameter space and away from the boundary of this area because perturbations to parameters would be neutral. This also implies that many genotypes correspond to the same phenotype. Therefore, these maps have the property of robustness but are also evolvable because they can neutrally explore phenotypic space as genetic variation can accumulate by mutation without changing the phenotype. This metaphor of the GP map has been used to study the evolution of RNA folding, protein structure, and gene-regulatory networks, for example (5). More recently, Nichol et al. (6) provide a more general definition for the GP map. These authors define a GP map as a relation that maps a genotype (along with a mutation operation) to a phenotype modulated by an environment. They further provide examples of such maps (phenotype landscapes, fitness landscapes, RNA secondary structures, gene-regulation, and neural networks for phenotypic plasticity) and discuss their general properties such as degeneracy, robustness, evolvability, and neutrality. By extending the definition such that the phenotype is modified by the environment, phenotypic plasticity can also be placed within the framework of GP maps. Additionally, this definition does not assume the GP map properties and includes more classical approaches as well. Therefore, we henceforward will use the definition given by Nichol et al. (6) and will refer to the conceptualisation by Alberch (4) particularly if required.

A very well studied GP map is the gene-regulatory network (GRN) model, particularly the one introduced by Wagner (7) to study gene duplications. The GRN represents a developmental process. The GRN GP map consists of $n$ transcription factors which can regulate each other's gene expression states (which are the phenotype), and the interaction between the genes is represented by a regulatory matrix (the genotype) containing weights. These weights are optimised by evolutionary simulations, where fitness is a function of gene expression levels (phenotype). Note that gene expression states in this model are usually discrete, that is, genes are either on or off. This is conceptually similar to the GP map described by Alberch (4) and has analogous properties. The parameters of the dynamical developmental system are the weights, and the gene expression levels are the steady states of this system. Various applications and variants of this model are reviewed in Spirov and Holloway (8). Particularly, the robustness to mutation, also known as genetic canalisation, has been studied in GRNs extensively. Wagner (9) suggests that robustness to mutation in GRNs evolves due to stabilising selection. Siegal and Bergman (10) however, suggest that GRNs can evolve to be robust in the absence of stabilising selection when there is selection for developmental stability. Ciliberti et al. (11) visualise different GRN genotypes as a genotype network, in which genotypes with a single mutation are connected to each other. They find that the genotype network can mostly be explored without a change in the phenotype, and particular genotype network topologies can be simultaneously robust and innovate. Rünneburger and Rouzic (12) challenge the generalisability of the evolution of robustness in GRN models when gene expression is quantitative rather than discrete. They model a GRN with sigmoid, continuous gene expression. Stabilising selection acts on few genes directly (selected genes), and the remaining genes are under indirect selection (only their regulatory function is selected). They find that, while the genes that are not directly selected always become canalised, selected genes are only canalised when there is selection for extreme levels of gene expression. This raises the further question of whether and how Wagner's model can be used to understand phenotypes that are quantitative rather than discrete. Furthermore, can this model be extended in a meaningful way to understand complex traits, which do not necessarily result from transcription regulation alone but from the downstream effects of regulatory genes? Is robustness to mutation still a feature of such a model?

GRN models have also been used to study the evolution of evolvability under conditions of ecological change, including local adaptation. Draghi and Wagner (13) use the GRN model to study the evolution of evolvability. They conclude that evolvability can evolve, and networks with greater positive network excitation are more evolvable but less robust under fluctuating selection. Kimbrell and Holt (14) use Wagner's model to understand local adaptation in a source-sink context. Optimal gene expression states are different in the source and sink. They find that adaptation to the sink happens in one or more steps and results from a breakdown in canalisation of the local adaptation GRN, that is, an increase in sensitivity to mutation. Kimbrell (15) extends this model and finds that sensitivity to mutation increases serially along a gradient landscape with four patches. Malcom (16) models adaptation to an environmental shift governed by a quantitative trait encoded by the sum of expression states of a boolean GRN. They find that smaller GRNs lead to a faster rate of adaptation. In Malcom (17), they extend this model to a metacommunity of two competing species of different network sizes and three patches. A local adaptation quantitative trait is represented by the sum of outputs of a Boolean GRN, and dispersal is a fixed rate. They find that for species with similar GRN sizes, the outcome of competition is determined by dispersal, and for comparable dispersal rates, the outcome of competition is determined by which species is able to adapt faster (those with networks of a smaller size). Finally, Melián et al. (18) suggest that including GRNs to model traits relevant to interspecific interactions can help us understand the stability and complexity of spatial communities like food webs.

#### Gene-regulatory network model for the evolution of dispersal and local adaptation during range expansion

The GRN assumed here is a modified version of the model by Wagner (7). However, as opposed to Wagner (7) and similar to Draghi and Whitlock (19), we assume that relevant traits (dispersal and local adaptation) are continuous and result from the downstream effects of a GRN. The GRN model by Wagner (7) assumes that a single developmental stage in a single cell type (tissue) is specified by equilibrium expression of $n$-transcription factors (either on or off). These $n$-transcription factors are represented by a discrete dynamical system and interact with each other cooperatively (switch-like response) during development. Such interactions are represented by regulatory matrices, which are the genotype of the individual. Thus, the GRN model maps a genotype (regulatory matrix) to a phenotype (gene expression levels) via a developmental process.

We introduce changes to the model by Wagner (7), which are similar to Draghi and Whitlock (19) and van Gestel and Weissing (20). Like van Gestel and Weissing (20), we visualise the model as having three layers, an input layer, a regulatory layer, and an output layer (Fig. 1). The input layer $\mathbf{x} = (x_1, ..., x_m)$ is a vector of upstream signals which may result from a previous developmental step if one is modelling a constitutive trait or a set of environmental cues to model phenotypic plasticity. Therefore, since we assume that dispersal and local adaptation are constitutive, we provide a fixed input $x_1 = 0.5$ to both the dispersal and local adaptation GRNs. The regulatory layer represents the network of n-transcription factors as modelled in Wagner (7). The normalised and transformed expression states of the regulatory genes after $I$ iterations are $\mathbf{S_I} = (S_1, ..., S_n)$. The genes can take values in the interval $[-1, 1]$ and gene expression of a single gene is a continuous sigmoid function (10, 19) of the inputs it receives. Each gene receives inputs from the input layer, itself, or other regulatory genes. An $m \times n$ matrix $\boldsymbol{U}$ connects the input layer to the regulatory layer. We call this the input matrix. Interactions between the genes, including autoregulatory interactions, are represented by an $n \times n$ matrix $\boldsymbol{W}$, termed the regulatory matrix. In a slightly different approach from Wagner (7), the properties of individual genes can also evolve. Genes can have slopes $\mathbf{r} = (r_1, ..., r_n)$ and thresholds $\theta = (\theta_1, ..., \theta_n)$ (20). A high slope would mean that gene expression as a function of the input is switch-like, and a shallow slope would mean that it is more gradual. When the input to the gene is greater than the threshold, then gene expression is positive, if the slope is also positive. The equation below describes the dynamic behaviour of the $i^{th}$ gene in the GRN.

$$S_{i,I+1} = \frac{2}{1 + \exp\left(-r_i(\sum_{j=1}^{j=m} U_{ji}x_j + \sum_{k=1}^{k=n} W_{ki}S_{k,I} - \theta_i)\right)} - 1 \tag{S5}$$

The network is iterated to obtain equilibrium gene expression, which can be a fixed point or a limit cycle. Limit cycles are considered to represent developmental instability (7) and individuals with such GRNs die. While oscillatory GRNs are possible in biological scenarios (especially those that involve timekeeping, for example, circadian rhythms, Novák and Tyson 21), the traits under consideration here, that is, dispersal and local adaptation, are characterized by a single value. Unless there is an even more complex mapping that converts the oscillatory expression states to a steady dispersal phenotype, the choice to select for developmental stability is reasonable, given that all individuals disperse at the same time in this model. However, one must be aware that this choice has consequences for evolved GRN properties, for example, the robustness of the GRN to mutations. Siegal and Bergman (10) find that selection for developmental stability leads to more robust GRNs. Espinosa-Soto (22) find that stabilising selection for particular gene expression states, including limit cycles, leads to the evolution of mutational robustness. Note that this makes any results related to the evolution of sensitivity in our model conservative.

We assume that genes in this GRN could be stochastically on or off before the beginning of the particular developmental state being modelled. Therefore, the initial gene expression states are drawn from a uniform distribution in the interval $[-1, 1]$. It is possible that the equilibrium gene expression could change depending on this initialisation. However, such networks are not highly frequent (23), and if the magnitude of the shift is high, then such a network might be eliminated from the metapopulation because of the fitness consequence of an incorrect dispersal decision or mismatch in trait optimum.

Similar to Draghi and Whitlock (19), we do not consider the vector of gene expression states as the phenotype. The phenotype is continuous and results from downstream effects of regulatory genes (for example, expression levels of structural genes or metabolic pathways). Further, we assume that the phenotype varies linearly with gene expression states. Hence, this model applies to cases where the processes downstream of regulatory gene expression are continuous and linear. However, the downstream effects of regulatory gene expression can be switch-like or non-linear as well (for example, the decision of a bacterium to sporulate in van Gestel and Weissing 20). This will not be considered in the present study. Therefore, the phenotypic value of a trait $(z)$ is given by linear combination of the gene expression levels at fixed point steady-state gene expression $\mathbf{S}^*$. $\boldsymbol{V}$ is a $1 \times n$ matrix containing weights connecting the regulatory layer to the output layer.

$$z = \sum_{i=1}^{n} V_i S_i^* \tag{S6}$$

We assume that two distinct GRNs encode dispersal ($z = d$) and local adaptation ($z = \tau_{opt}$). The gene expression states are iterated for 20 iterations $I$ and then the continuous phenotype is calculated.

##### Calculation of the sensitivity to mutation in the GRN model

In the gene-regulatory network model we assume that the loci interact non-additively to output a given trait. Therefore, how mutation changes a phenotype can evolve. This means that under conditions of strong stabilising selection on the trait under consideration, the GRNs can become more robust (less sensitive) to mutation (for example 9). The sensitivity to mutation can also increase under directional selection for the optimum of a trait to change (for example, Kimbrell and Holt 14). To quantify the sensitivity to mutation of a trait (here, dispersal or local adaptation), we use an approach similar to Kimbrell and Holt (14), Kimbrell (15) and Draghi and Whitlock (19). The sensitivity to mutation of a trait is calculated by sampling individuals from a specified region in the landscape (range core, range front or a given patch cross-section) at a given time, and for each individual introducing 100 haploid perturbations drawn from $N(0, 0.1)$ at one randomly selected locus of the GRN for that trait. The root mean squared difference between the perturbed, and the unperturbed phenotype over all the haploid perturbations and individuals is the sensitivity to mutation. To put it simply, it is a metric of how much the trait changes at the population level when a haploid mutation is introduced to one of the loci encoding it. Note that sensitivity can change only in the case of the GRN model but is fixed in the single locus and multilocus models because the latter genetic architectures are additive.

### S1 Supplementary tables

| Model Parameter/-Variable | Description | Values |
|---|---|---|
| $N_{x,y,t}$ | Population density in the patch $x, y$ at time $t$ | dynamical |
| $\lambda_0$ | Intrinsic growth rate in Beverton-Holt model | 2 |
| $\alpha$ | Intra-specific competition coefficient in Beverton-Holt model | 0.01 |
| $\mu$ | Dispersal cost | 0.01, 0.1, 0.3 |
| $\epsilon$ | Local patch extinction probability | 0, 0.1 |
| $m_{min}$ | Mutation rate at equilibrium | 0.0001 |
| $m_{max}$ | Mutation rate at the beginning | 0.1 for GRN, $m_{max} = m_{min}$ for others |
| $\sigma_m$ | Effect size (standard deviation) of mutations | 0.1 |
| $\mathbf{x}$ | Vector of inputs to the GRN | in simulation |
| $\mathbf{S}_I$ | Vector of expression states of regulatory genes at iteration $I$ | in simulation |
| $\boldsymbol{U}$ | $m \times n$ matrix with each element $U_{ji}$ representing the connection between the input $j$ and gene $i$ | evolves, initialised from a normal distribution with sd = 1 |
| $\boldsymbol{W}$ | $n \times n$ matrix with each element $W_{ki}$ representing the connection between the gene $k$ and gene $i$ | evolves, initialised from a normal distribution with sd = 1 |
| $\boldsymbol{V}$ | $1 \times n$ matrix with each element $V_i$ representing the connection between the gene $k$ and the output | evolves, initialised from a normal distribution with sd = 1 |
| $\boldsymbol{\theta}$ | Thresholds of regulatory genes | evolves, initialised from a normal distribution with sd = 1 |
| $\mathbf{r}$ | Slopes of regulatory genes | evolves, initialised from a normal distribution with sd = 1 |
| $\tau(x)$ | Value of environmental variable at position x | calculated from Eq. S3 |
| $\tau_0$ | Value of environmental variable in range core | 0.1 |
| $b$ | Slope of environmental gradient | 0.02, 0.04 |
| $\tau_{opt}$ | Niche optimum | evolves, encoded either by a GRN or single locus |
| $\omega$ | Niche width | 1 |

Table S1: Model Parameters/Variables

### Supplementary figures

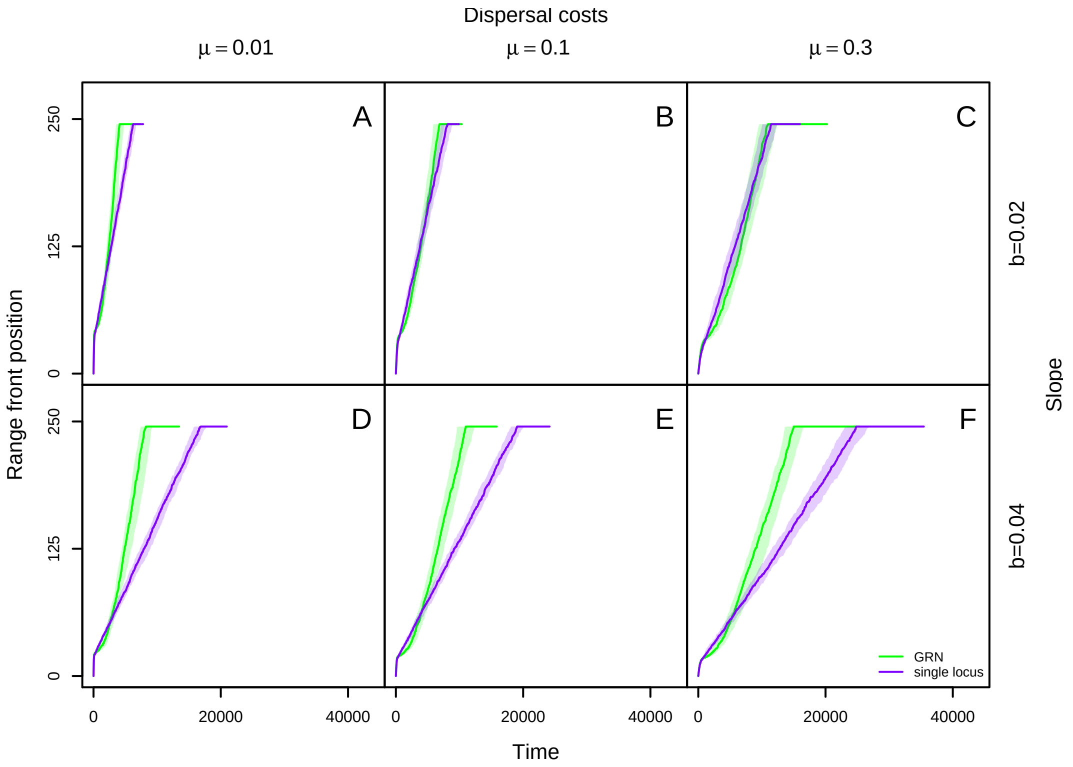


Figure S1: Effect of dispersal costs $\mu$ and slope of environmental gradient $b$ on range front position as a function of time since the beginning of range expansion when there are no random patch extinctions ($\epsilon = 0$)—single locus and GRN models. The solid lines indicate median position of the range front (defined as the occupied patch farthest from the range core) at every 50 time steps over 100 range expansions and the shaded region represents the quartiles. From left to right, dispersal costs increase and from top to down slope of the environmental gradient increases. The general pattern of accelerated range expansions in the GRN model but not in the single locus model is robust to changes in dispersal costs and the slope of the environmental gradient. Fixed parameters: female fecundity $\lambda_0 = 2$, intra-specific competition coefficient $\alpha = 0.01$, niche width $\omega = 1$, mutation rate during range expansion $m_{min} = 0.0001$, number of genes per GRN $n = 3$.

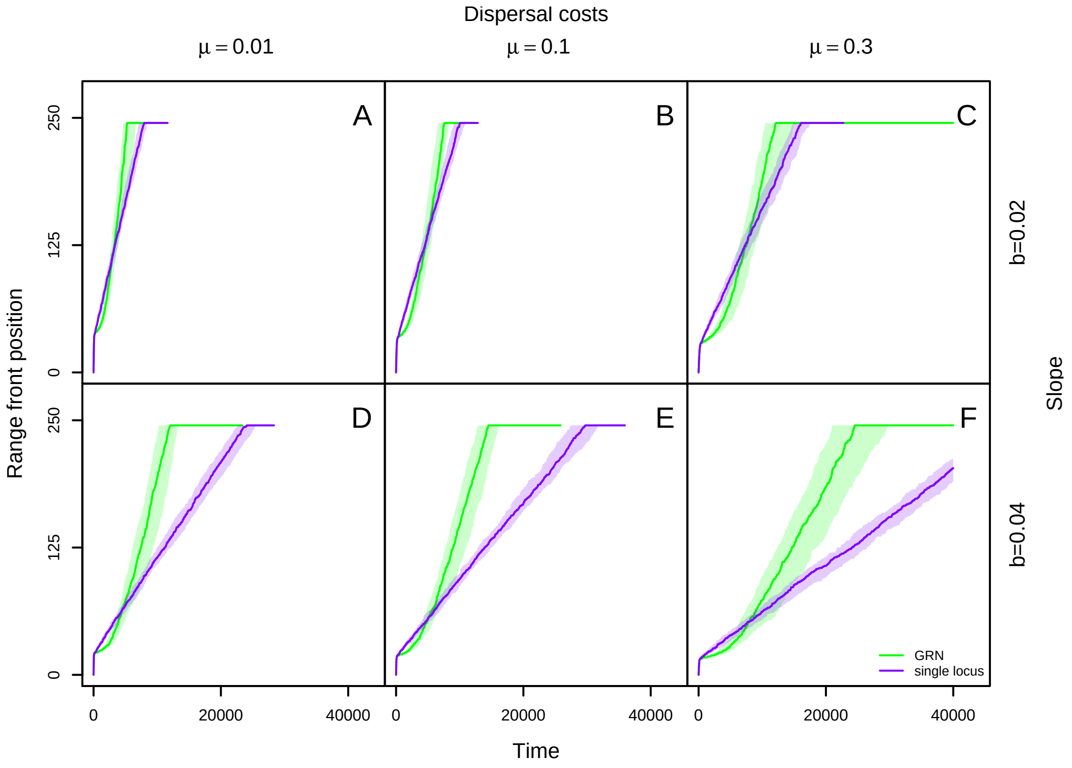


Figure S2: Effect of dispersal costs $\mu$ and slope of environmental gradient $b$ on range front position as a function of time since the beginning of range expansion when there are random patch extinctions with a probability $\epsilon = 0.1$—single locus and GRN models. The solid lines indicate median position of the range front (defined as the occupied patch farthest from the range core) at every 50 time steps over 100 range expansions and the shaded region represents the quartiles. From left to right, dispersal costs increase and from top to down slope of the environmental gradient increases. The general pattern of accelerated range expansions in the GRN model but not in the single locus model is robust to changes in dispersal costs, the slope of the environmental gradient and extinction probability. Fixed parameters: female fecundity $\lambda_0 = 2$, intra-specific competition coefficient $\alpha = 0.01$, niche width $\omega = 1$, mutation rate during range expansion $m_{min} = 0.0001$, number of genes per GRN $n = 3$.

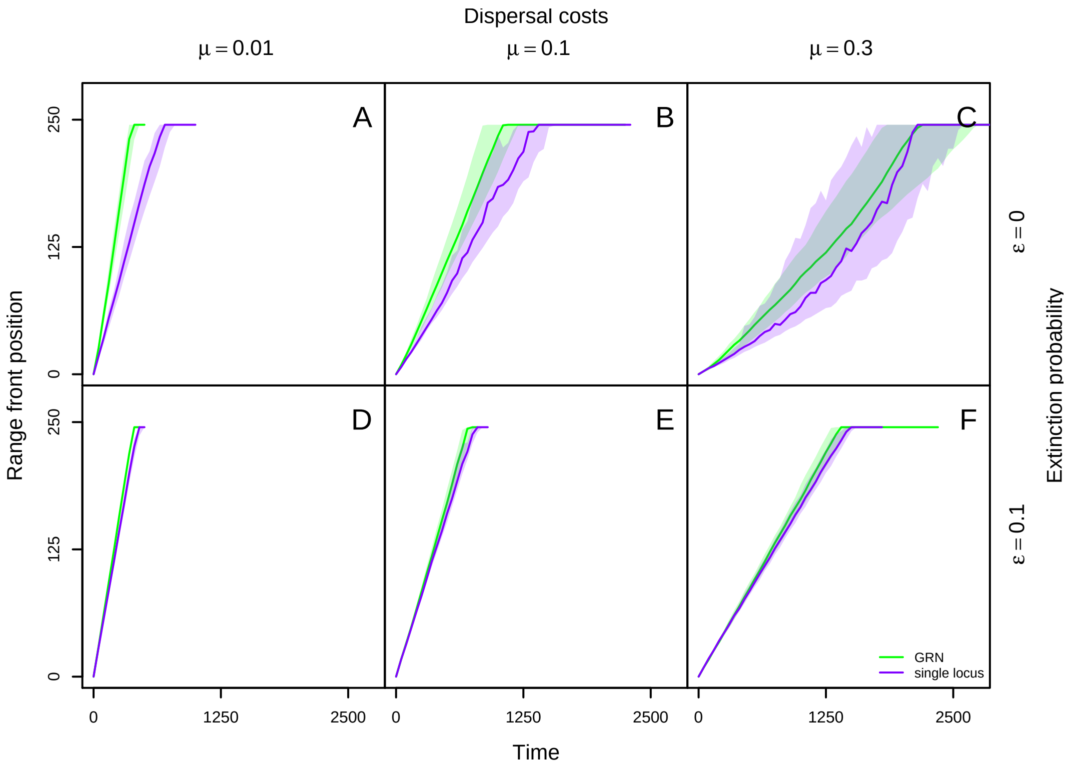


Figure S3: Range front position as a function of time in the absence of an external environmental gradient, with only dispersal evolving—single locus and GRN models. The solid lines indicate median position of the range front (defined as the occupied patch farthest from the range core) at every 50 time steps over 100 range expansions and the shaded region represents the quartiles. From left to right, dispersal costs increase and from top to down extinction probability increases. Fixed parameters: female fecundity $\lambda_0 = 2$, intra-specific competition coefficient $\alpha = 0.01$, mutation rate during range expansion $m_{min} = 0.0001$, number of genes per GRN $n = 3$.

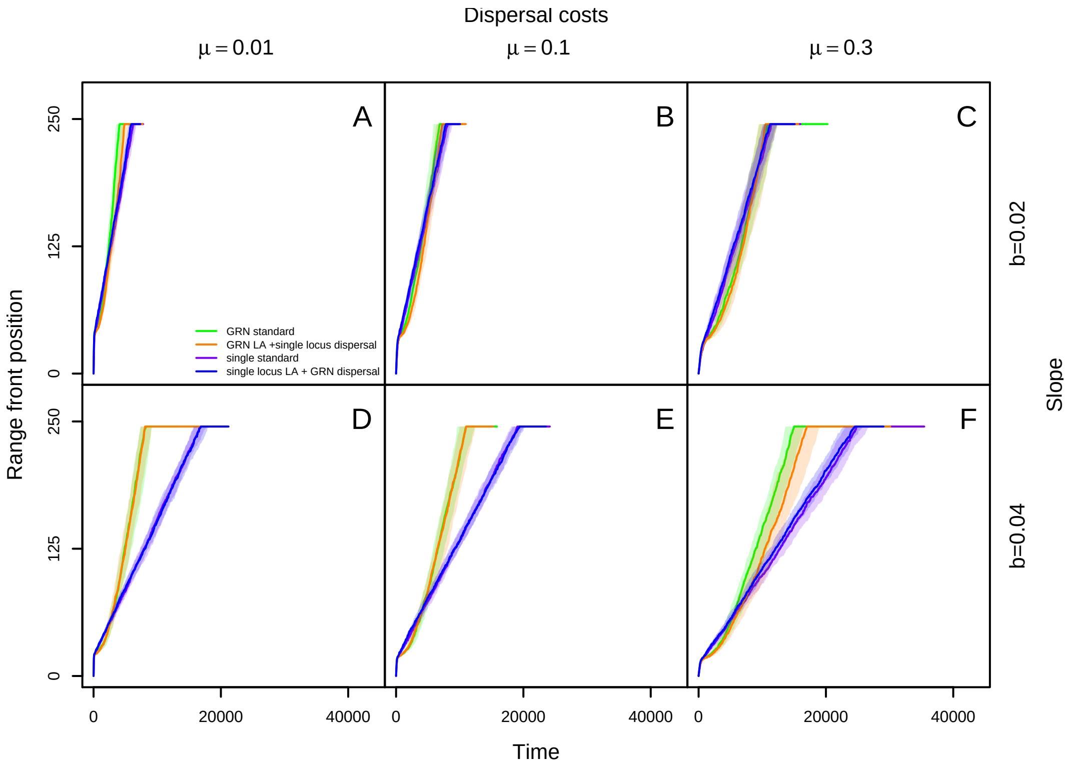


Figure S4: Effect of genetic architecture of dispersal on range front position as a function of time since the beginning of range expansion when there are no patch extinctions ($\epsilon = 0$)—GRN encodes both dispersal and local adaptation (GRN model), single locus each encodes dispersal and local adaptation (single locus model), GRN encodes local adaptation and a single locus encodes dispersal (GRN LA+single locus dispersal model) and single locus encodes local adaptation and a GRN encodes dispersal (single locus LA+GRN dispersal model). The solid lines indicate median position of the range front (defined as the occupied patch farthest from the range core) at every 50 time steps over 100 range expansions and the shaded region represents the quartiles. From left to right, dispersal costs increase and from top to down the slope of the environmental gradient increases. Range expansions are accelerated if local adaptation is encoded by a GRN irrespective of the genetic architecture of dispersal for different dispersal costs and slope of external environmental gradient. Fixed parameters: female fecundity $\lambda_0 = 2$, intra-specific competition coefficient $\alpha = 0.01$, niche width $\omega = 1$, mutation rate during range expansion $m_{min} = 0.0001$, number of genes per GRN $n = 3$.

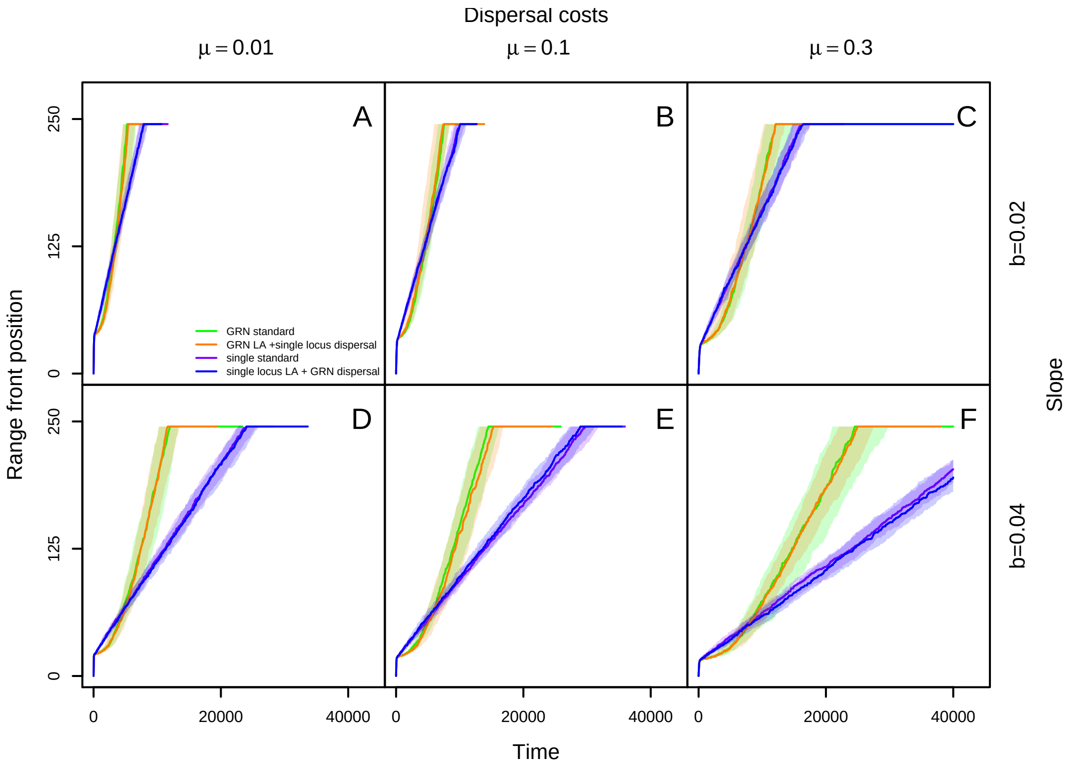


Figure S5: Effect of genetic architecture of dispersal on range front position as a function of time since the beginning of range expansion when there are patch extinctions with probability $\epsilon = 0.1$—GRN encodes both dispersal and local adaptation (GRN model), single locus each encodes dispersal and local adaptation (single locus model), GRN encodes local adaptation and a single locus encodes dispersal (GRN LA+single locus dispersal model) and single locus encodes local adaptation and a GRN encodes dispersal (single locus LA+GRN dispersal model). The solid lines indicate median position of the range front (defined as the occupied patch farthest from the range core) at every 50 time steps over 100 range expansions and the shaded region represents the quartiles. From left to right, dispersal costs increase and from top to down the slope of the environmental gradient increases. Range expansions are accelerated if local adaptation is encoded by a GRN irrespective of the genetic architecture of dispersal for different dispersal costs and slope of external environmental gradient, even when there are patch extinctions. Fixed parameters: female fecundity $\lambda_0 = 2$, intra-specific competition coefficient $\alpha = 0.01$, niche width $\omega = 1$, mutation rate during range expansion $m_{min} = 0.0001$, number of genes per GRN $n = 3$.

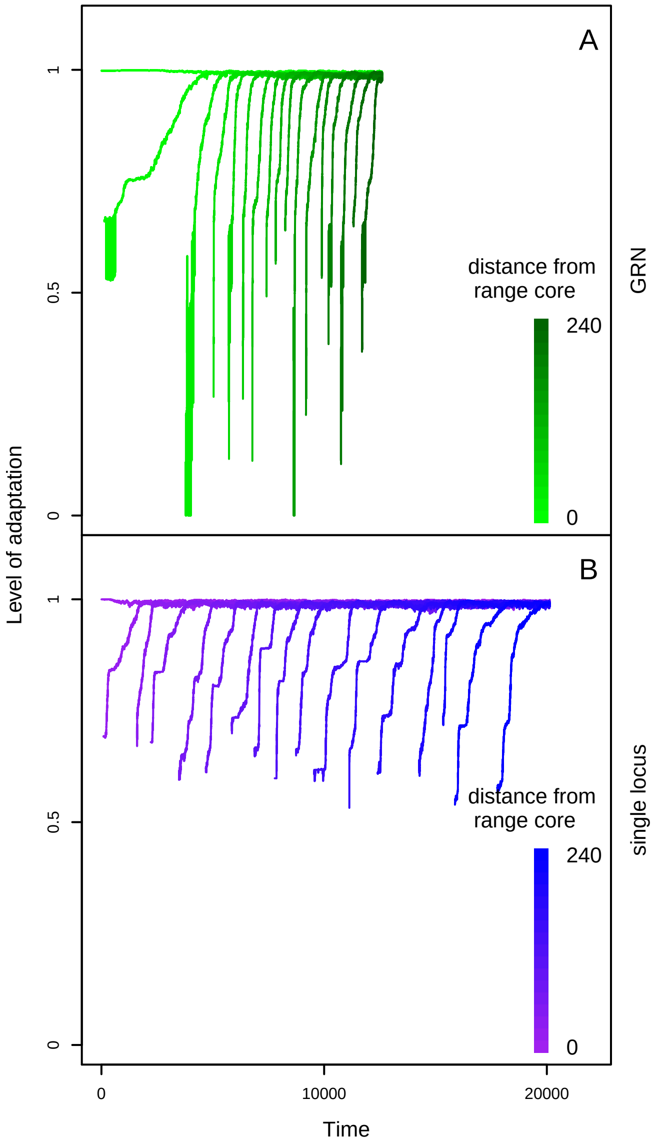


Figure S6: Example dynamics of local adaptation to a given environment—GRN and single locus models. The level of adaptation $s$, as a function of time since the beginning of range expansion shown for every $15^{th}$ patch from the range core for a single range expansion. A: GRN model B: single locus model. Darker colours indicate patches farther away from the range core. The expanding population adapts more quickly to patches farther from the range core in the GRN model but not in the single locus model. Focal scenario parameters: slope of gradient $b = 0.04$, fecundity of the females $\lambda_0 = 2$, intra-specific competition coefficient $\alpha = 0.01$, mutation rate during range expansions $m_{min} = 0.0001$, dispersal cost $\mu = 0.1$, extinction probability $\epsilon = 0$, niche width $\omega = 1$, number of genes per GRN $n = 3$.

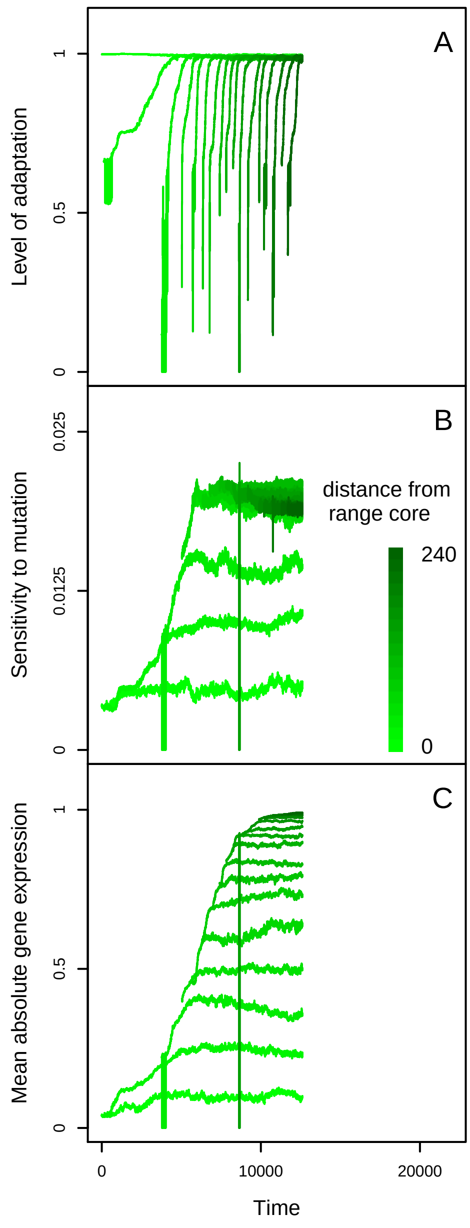


Figure S7: Example dynamics for the level of adaptation of the expanding population, sensitivity to mutation and mean absolute gene expression of the local adaptation trait—GRN model. A: level of adaptation to an environment as a function of time since the beginning of range expansion. B: Sensitivity to mutation of the local adaptation trait as a function of time since the beginning of range expansion. C: Averaged absolute gene expression levels over the three genes governing the local adaptation trait as a function of time since the beginning of range expansion. In all the plots, darker shades of green indicate that the calculation is for an environment farther away from the range core. Focal scenario parameters: slope of gradient $b = 0.04$, fecundity of the females $\lambda_0 = 2$, intra-specific competition coefficient $\alpha = 0.01$, mutation rate during range expansions $m_{min} = 0.0001$, dispersal cost $\mu = 0.1$, extinction probability $\epsilon = 0$, niche width $\omega = 1$, number of genes per GRN $n = 3$.

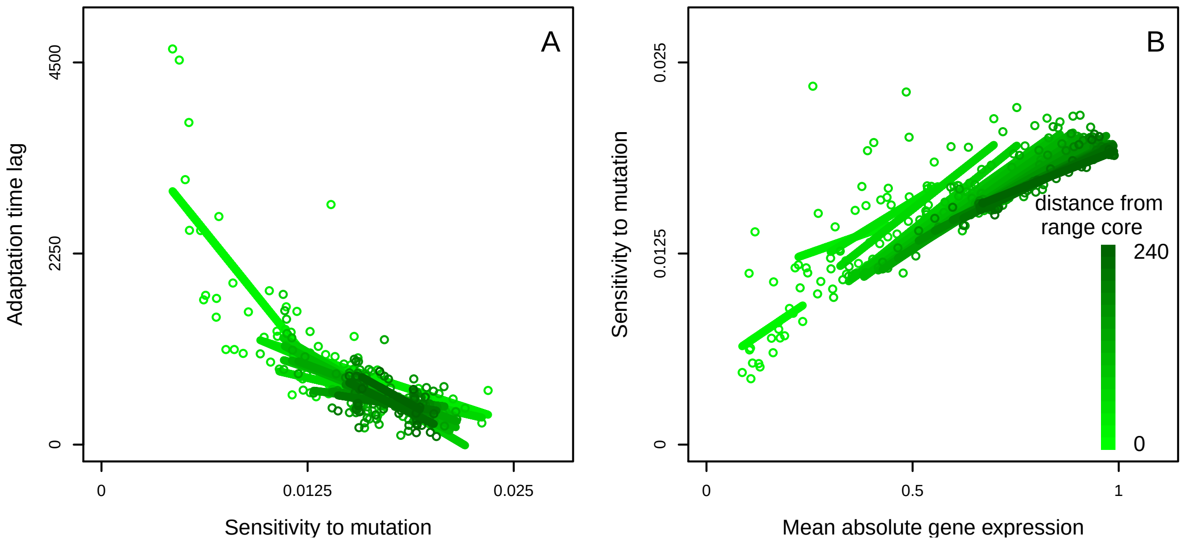


Figure S8: Relationship between the adaptation time lag of the expanding population and the sensitivity to mutation of the local adaptation trait, and between the sensitivity to mutation and mean absolute gene expression in the local adaptation GRN for a given environment—GRN model for dispersal evolution and local adaptation. A: Adaptation time lag as a function of the sensitivity to mutation of the local adaptation trait, for 10 replicates or 20 range expansions, colour coded by the distance from the landscape core. The lines of the same colour are fit by simple linear regression. B: Sensitivity to mutation of the local adaptation trait as a function of mean absolute gene expression of the local adaptation GRN, over 10 replicates or 20 range expansions, colour coded by the distance from the landscape core. The lines of the same colour are fit by simple linear regression. For a given patch cross section, expanding populations that are more sensitive to mutations adapt more quickly and are associated with extremes of gene expression. Focal scenario parameters: slope of gradient $b = 0.04$, fecundity of the females $\lambda_0 = 2$, intra-specific competition coefficient $\alpha = 0.01$, mutation rate during range expansions $m_{min} = 0.0001$, dispersal cost $\mu = 0.1$, extinction probability $\epsilon = 0$, niche width $\omega = 1$, number of genes per GRN $n = 3$.

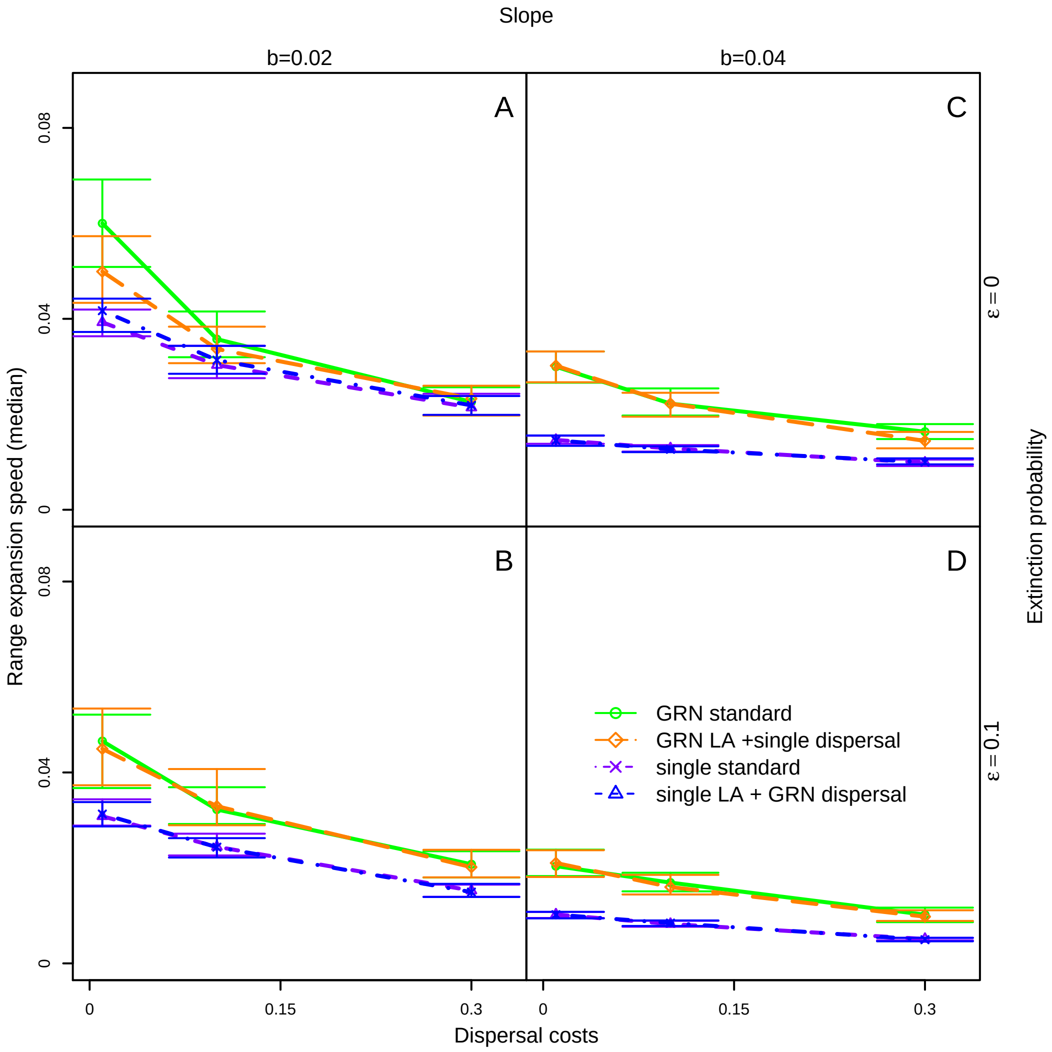


Figure S9: Effect of slope of environmental gradient, dispersal costs, extinction probability and genetic architecture of dispersal on the median of the speed of range expansion—GRN encodes both dispersal and local adaptation (GRN model), single locus each encodes dispersal and local adaptation (single locus model), GRN encodes local adaptation and a single locus encodes dispersal (GRN LA+single locus dispersal model) and single locus encodes local adaptation and a GRN encodes dispersal (single locus LA+GRN dispersal model). The median of expansion speed over 50 replicates, or 100 range expansions as a function of dispersal costs. From left to right, slope increases, from to top to bottom, extinction probability increases. Fixed parameters: female fecundity $\lambda_0 = 2$, intra-specific competition coefficient $\alpha = 0.01$, mutation rate during range expansion $m_{min} = 0.0001$, number of genes per GRN $n = 3$.

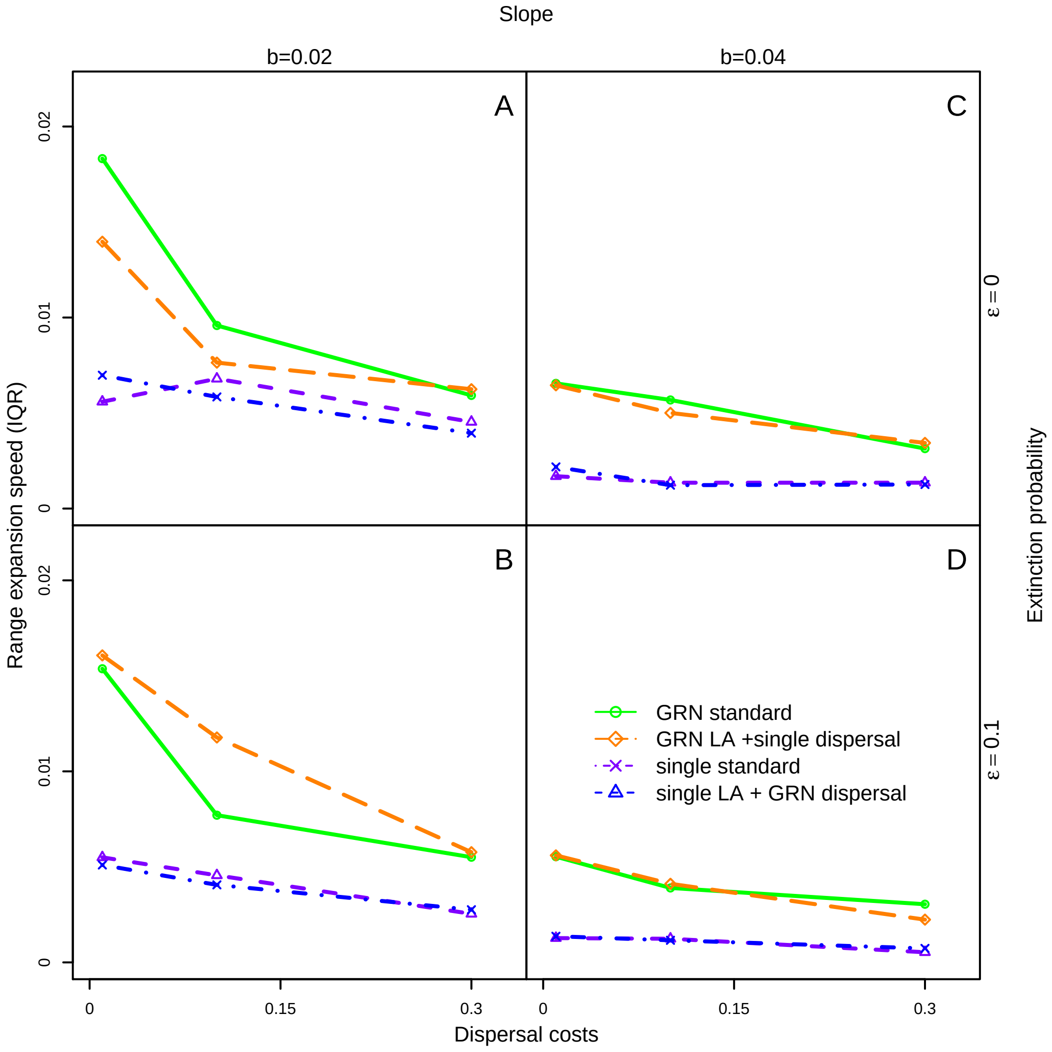


Figure S10: Effect of slope of environmental gradient dispersal costs, extinction probability and genetic architecture of dispersal on the interquartile range of the speed of range expansion—GRN encodes both dispersal and local adaptation (GRN model), single locus each encodes dispersal and local adaptation (single locus model), GRN encodes local adaptation and a single locus encodes dispersal (GRN LA+single locus dispersal model) and single locus encodes local adaptation and a GRN encodes dispersal (single locus LA+GRN dispersal model). The interquartile range of expansion speed over 50 replicates, or 100 range expansions as a function of dispersal costs. From left to right, slope increases, from to top to bottom, extinction probability increases. Fixed parameters: female fecundity $\lambda_0 = 2$, intra-specific competition coefficient $\alpha = 0.01$, mutation rate during range expansion $m_{min} = 0.0001$, number of genes per GRN $n = 3$.

### Supplementary References

[1] Bonte, D., Van Dyck, H., Bullock, J. M., Coulon, A., Delgado, M., Gibbs, M., Lehouck, V., Matthysen, E., Mustin, K., Saastamoinen, M., Schtickzelle, N., Stevens, V. M., Vandewoestijne, S., Baguette, M., Barton, K., Benton, T. G., Chaput-Bardy, A., Clobert, J., Dytham, C., Hovestadt, T., Meier, C. M., Palmer, S. C. F., Turlure, C., & Travis, J. M. J. (2012) *Biol. Rev. Biol. Proc. Camb. Philos. Soc.* **87**, 290–312.

[2] Beverton, R. J. H. & Holt, S. J. (1957) *On the dynamics of exploited fish populations* (Chapman & Hall, London).

[3] Poethke, H. J. & Hovestadt, T. (2002) *Proc. R. Soc. Lond. B Biol. Sci.* **269**, 637–645.

[4] Alberch, P. (1991) *Genetica* **84**, 5–11.

[5] Pigliucci, M. (2010) *Philos. Trans. R. Soc. B-Biol. Sci.* **365**, 557–566.

[6] Nichol, D., Robertson-Tessi, M., Anderson, A. R. A., & Jeavons, P. (2019) *J. R. Soc. Interface* **16**, 20190332.

[7] Wagner, A. (1994) *Proc. Natl. Acad. Sci.* **91**, 4387–4391.

[8] Spirov, A. & Holloway, D. (2013) *Methods* **62**, 39–55.

[9] Wagner, A. (1996) *Evolution* **50**, 1008.

[10] Siegal, M. L. & Bergman, A. (2002) *Proc. Natl. Acad. Sci.* **99**, 10528–10532.

[11] Ciliberti, S., Martin, O. C., & Wagner, A. (2007) *Proc. Natl. Acad. Sci.* **104**, 13591–13596.

[12] Rünneburger, E. & Rouzic, A. L. (2016) *BMC Evol. Biol.* **16**.

[13] Draghi, J. & Wagner, G. P. (2009) *J. Evol. Biol.* **22**, 599–611.

[14] Kimbrell, T. & Holt, R. (2007) *Am. Nat.* **169**, 370–382.

[15] Kimbrell, T. (2010) *Evol. Ecol.* **24**, 891–909.

[16] Malcom, J. W. (2011) *PLoS ONE* **6**, e14747.

[17] Malcom, J. W. (2011) *PLoS ONE* **6**, e21541.

[18] Melián, C. J., Matthews, B., de Andreazzi, C. S., Rodríguez, J. P., Harmon, L. J., & Fortuna, M. A. (2018) *Trends Ecol. Evol.* **33**, 504–512.

[19] Draghi, J. A. & Whitlock, M. C. (2012) *Evolution* **66**, 2891–2902.

[20] van Gestel, J. & Weissing, F. J. (2016) *Sci. Rep.* **6**.

[21] Novák, B. & Tyson, J. J. (2008) *Nat. Rev. Mol. Cell Biol.* **9**, 981–991.

[22] Espinosa-Soto, C. (2016) *J. Evol. Biol.* **29**, 2321–2333.

[23] Nido, G. S., Williams, J. M., & Benuskova, L. (2012). Bistable properties of a memory-related gene regulatory network.
